## Supplemental Results for "Germline polymorphisms in the immunoglobulin kappa and lambda loci explain variation in the expressed light chain antibody repertoire"

**This PDF file includes:**

Supporting text
Figures S1 to S21
SI References

**Supporting Information Text**

**Identification of SVs, SNVs, and gene alleles in IGK and IGL using long-read sequencing**

To genotype variants in IGK and IGL, we used probe-based targeted capture long-read single molecule real-time (SMRT) sequencing [1] of the IGK proximal and distal regions [2], and the IGL locus [3]. IGK was sequenced to a coverage of 65.9X on average, with a mean of 1,858,094 diploid base pairs assembled per individual and mean assembly accuracy of 99.96%. IGL was sequenced to a coverage of 41.3X on average, with a mean of 1,903,810 diploid base pairs assembled per individual and mean assembly accuracy of 99.93% (**Supplementary Table S1, Supplementary Fig. 1**).

One of the primary objectives of this study was to develop a high-confidence set of genetic variants in IGK and IGL to enable downstream genetic association analysis. We previously described common SVs in IGK [2] and IGL [3], including polymorphisms associated with gene conversion in IGK, a large inversion in IGK, a deletion involving IGKV1-NL1, a deletion involving IGLV5-39, and a deletion involving IGLV3-16, IGLV2-18, IGLV3-19, IGLV3-21, IGLV3-22, and IGLV2-23 (termed IGLV2-18) (**Supplementary Fig. 2A-B**).

In addition to these SVs, we identified a haplotype wherein the entire IGKV distal region is deleted, which was previously reported [4,5], removing 23 functional IGKV genes. Nine individuals with this deletion haplotype exhibited ~2-3-fold higher sequencing coverage over the proximal relative to distal region (**Supplementary Fig. 4**), suggestive of hemizygosity. An additional individual lacked both HiFi reads and assemblies mapped to the IGK distal region, despite having 87X coverage and diploid assemblies for the proximal region (**Supplementary Fig. 4**), indicating homozygous absence of the IGKV distal region in this sample. SV allele frequencies for samples in this cohort are summarized in **Supplemental Table S2**.

SNVs within segmental duplications are difficult to characterize. We previously demonstrated that SNV callsets from haplotype-resolved IGK [2] and IGL [3] assemblies derived from long-read SMRT sequencing outperform short-read derived variant calls in phase 3 [6] of the 1KGP. We identified variants for each individual and generated a callset of common SNVs, defined as those with a minor allele frequency (MAF) ≥ 0.05, totaling 2,792 and 5,198 common SNVs in IGK and IGL, respectively (**Supplementary Fig. 2C**). Comparison of these SNVs with those in the dbSNP “common” (MAF ≥ 0.01) catalog revealed substantial differences; 59.3% and 38.8% of common SNVs in IGK and IGL, respectively, were absent from this dbSNP catalog (**Supplementary Fig. 2D-E**). These data indicate that dbSNP lacks accurate genotype information for about half of the common variants in IGK and about a third of the common variants in IGL.

While the majority of common SNVs were intergenic (IGK, 95%; IGL, 97%), SNVs were identified within features of V and J genes, including coding exons, V gene introns, and RSS heptamers, spacers, and nonamers (**Supplementary Fig. 5**). 34 of 47 IGKV genes (72%) and 29 of 39 IGLV genes (74%) harbored at least one common SNV within a gene feature.

Analysis of AIRR-seq data critically relies on assignment of AIRR-seq reads to specific IG gene alleles, which are typically identified from a germline allele database. For a given individual, accurate AIRR-seq analysis requires inclusion of all of the individual’s alleles in the germline database to permit accurate assignment of reads to gene alleles; these assignments are used for analyzing a variety of Ab repertoire features, including gene usage and somatic hypermutation. We previously demonstrated that there is significant variation among the IGK [2] and IGL [3] gene alleles in the human population, and many of these alleles are not documented in the commonly used germline alleles database called the ImMunoGeneTics Information System (IMGT; imgt.org). With haplotype-resolved IG assemblies in-hand, one can use a personalized germline allele database for AIRR-seq analysis of the individual.

To annotate germline IGK and IGL gene alleles in this cohort, we used haplotype-resolved assemblies and identified both documented and undocumented (novel) alleles, defined as alleles absent from IMGT (https://www.imgt.org). In total, we identified 160 and 145 high-confidence novel IGKV and IGLV alleles, respectively, defined as alleles with exact matches to ≥10 HiFi reads that mapped to the position of the allele sequence in the assembly (**Supplementary Fig. 2F-I**; **Supplementary Table S3**, **Supplementary Table S4**). Among all IGKV and IGLV alleles, 67.4% and 62.8% were not documented in IMGT (**Supplementary Fig. 6**). Among the novel alleles, 81 IGKV and 70 IGLV alleles were identified in more than 1 individual (**Supplementary Fig. 2H-I**). The majority of novel alleles resulted in non-synonymous substitutions (as compared to the closest-matching allele in IMGT), with 7 IGKV and 7 IGLV novel alleles encoding premature STOP codons (**Supplementary Fig. 2H-I**). We noted that 44 novel IGKV alleles were also identified in our previous survey of IGKV alleles using samples collected as part of the 1KGP [2]. To access and explore curated genetic resources from this dataset further see Peres et al. (in prep).

**IGK and IGL repertoire-wide gene usage profiles are more highly correlated in individuals carrying shared genotypes**

Monozygotic twin studies have shown that gene usage frequencies in genetically identical individuals correlated to a greater degree than in unrelated individuals [7–9]. For IGH, we extended this observation at the population level and demonstrated that repertoire-wide gene usage profiles are more highly correlated in individuals carrying shared genotypes at guQTL SNVs [10]. To assess this in IGK and IGL, we used the same approach by estimating allele sharing distance (ASD) [11,12] in our cohort across IGK or IGL, and comparing the gene usage correlations between groups of individuals with higher and lower ASDs. Repertoire-wide gene usage correlations between samples were calculated using the Pearson's Correlation coefficient. Using all guQTL variants for each gene, individuals with the most overlapping guQTL genotypes (low ASD) had a higher mean gene usage correlation than those in the group with the highest ASD scores for both IGK (0.983 vs. 0.966; KS test $p$ = 4.4e-141) and IGL (0.925 vs. 0.900 ; KS test $p$ = 7.1e-03) (**Supplementary Fig. 10**). These results indicated that genetic background makes a contribution to the overall gene usage composition of the repertoire, and expand on observations from twin studies [7–9] by demonstrating that heritable components of the light chain repertoire can be directly linked to germline variants in light chain loci.

**Variants associated with IGL gene usage variation are enriched in regulatory regions**

Large-scale studies utilizing expression, epigenomic, and variant datasets associated with diseases or traits have uncovered non-coding variants within regulatory elements linked to specific phenotypes [13–16]. In the context of V(D)J recombination, RSS are recognized by RAG1/RAG2 proteins to direct double-strand DNA breaks and initiate somatic recombination [17]. This process is regulated by cis-elements that interact with chromatin-binding proteins which collectively determine the probability of somatic recombination for a given gene [18–22]. Given that non-coding variants associated with IGH gene usage variation are enriched in regulatory regions, we hypothesized that non-coding variants might impact light-chain gene usage by affecting regulatory elements.

We tested for enrichment of candidate transcription factor binding sites (TFBS, ENCODE3 Transcription Factor ChIP-seq dataset) in IGL (302 regulatory elements) and IGK (129 regulatory elements) overlapping lead guQTL variants in IGL (97 lead SNVs) and IGK (125 lead SNVs) versus the remainder of common SNVs for each locus, which included 1017 and 2310 non-lead variants in IGL and IGK, respectively. We found that lead IGL guQTLs were enriched with TFBS for multiple factors, including SRF, CBX1, TRIM28, TAL1, EGR1, and HNRNPUL1 (**Supplementary Fig. 12**). The guQTL variants overlapping these TFBS were associated with usage of IGLV3-10, IGLV2-8, IGLV3-9, IGLV3-27, IGLV2-11, and IGLV9-49 (**Supplementary Table S8**). In contrast with IGL, there was not a significant enrichment of lead IGK guQTL variants within TFBS.

The lead variant associated with the usage of IGLV3-10, IGLV2-8, and IGLV3-9 (**Supplementary Fig.12**) overlaps an EGR1 site in addition to >10 distinct annotated TFBS including SMC3, YY1, ETS1, ATF1, CTCF, and RAD21. Similarly, the lead variant associated with IGLV9-49 usage overlaps CBX1 and TRIM28 sites in additional to sites for >10 additional TF’s, among which were SMC3, CREB1, MYC, JUNB, STAT5A, CTCF, and RAD21 (**Supplementary Fig. 13**). We noted that the IGLV9-49 lead variant is 50 bp upstream of the IGLV9-49 5’ UTR.

These data suggest that a subset of IGL guQTLs are within cis-elements that may regulate V(D)J recombination and therefore gene usage in the peripheral antibody repertoire.

**Genetic variants disrupt biases in differential usage of IGKV proximal and distal paralogs**

IGKV proximal and distal gene paralogs vary substantially with respect to distance from the IGKJ region along the linear genome, with at least 1.0 Mbp separating distal V genes from J segments [23]. To date, there has not been an empirical evaluation of IGKV proximal versus distal gene usage. Historically, usage of IGKV genes has been difficult to assess due to high sequence similarity between paralogs [24,25]. With paired AIRR-seq and germline alleles available for the first time at population-scale, we determined the individual usage of 13 IGKV paralog pairs and observed higher usage of the proximal gene in 11 cases, with the exception of IGKV2-29/2-29, for which usage was not significantly different, and IGKV1-13/1D-13, for which distal paralog usage was higher (paired t-tests, *P* value < 1.0e-10) (**Supplementary Fig. 14A**).

IGKV1-13 was used at a lower frequency than IGKV1D-13 in all but one individual. Notably, all IGKV1-13 alleles have a non-canonical heptamer (CATAGTG) (**Supplementary Fig. 14B**), and IGKV1-13*01, the most frequent allele (frequency 73%, **Supplementary Fig. 14C**) encodes a premature STOP codon (**Supplementary Fig. 15**). Therefore, IGKV1-13 resembles a pseudogene, with both regulatory and coding loss-of-function features across haplotypes. IGKV1D-13 sequences, in contrast, included an allele (IGKV1D-13*02) with a canonical heptamer (CACAGTG) identified at a frequency of 23.6% (**Supplementary Fig. 14C**). The lead IGKV1D-13 guQTL tagged this heptamer variation, as all individuals in the higher-usage C/C genotype group were also IGKV1D-13*02/IGKV1D-13*02, whereas 69 of 70 individuals in the T/T genotype group did not carry the *02 allele (**Supplementary Fig. 14D**), implicating heptamer variation as a mechanism underlying usage variation.

IGKV2-29 was used at a lower frequency than IGKV2D-29 in 114 out of 170 individuals (67.1%). This inter-individual variation in IGKV2-29 versus IGKV2D-29 usage was explained by the lead IGKV2-29 guQTL, with 109 out of 110 individuals homozygous for the reference allele having higher IGKV2D-29 usage and 15 out of 15 individuals homozygous for the alternate allele having higher IGKV2-29 usage (**Supplementary Fig. 14F**). Mechanistically, the lead IGKV2-29 guQTL results in a premature STOP codon in the *01, *01_N1, and *01_N2 alleles (**Fig. 2B**), which are the only IGKV2-29 alleles carried by the 110 individuals homozygous for the guQTL reference allele.

**Supplementary Figures**

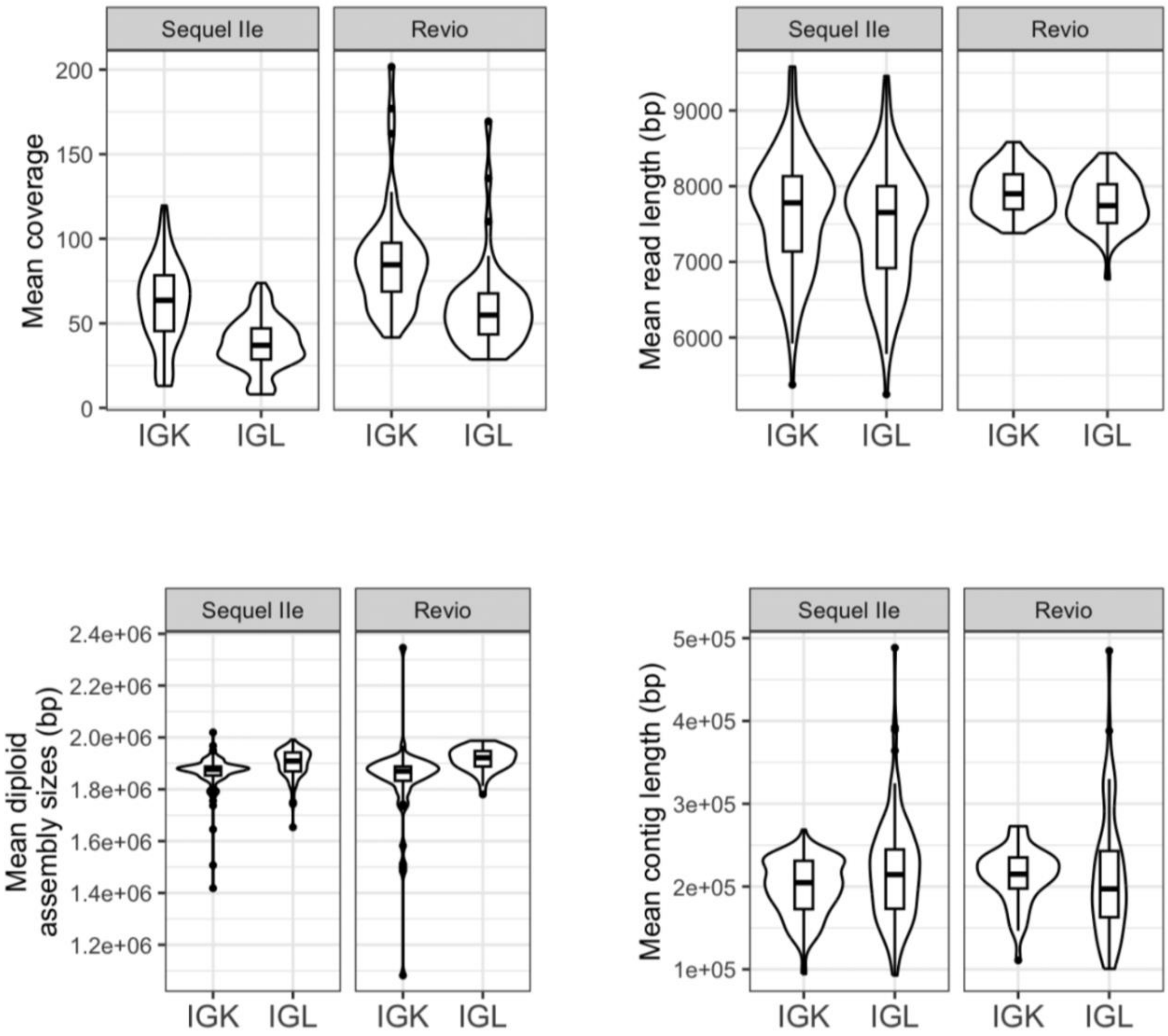


**Figure S1. PacBio sequencing and assembly statistics.**
For both IGK and IGL, the statistics described include (a) locus coverage, (b) read length, (c) assembly lengths, and (d) contig lengths. Each statistic is separated by PacBio platforms. The number of sequencing runs on the Sequel IIe and Revio platforms is 134 and 43, respectively. Boxplots display the median, 25th percentile, 75th percentile, and whiskers that extend up to 1.5 times the inter-quartile range (IQR) from the respective percentiles. Any data points outside the whiskers are also plotted.

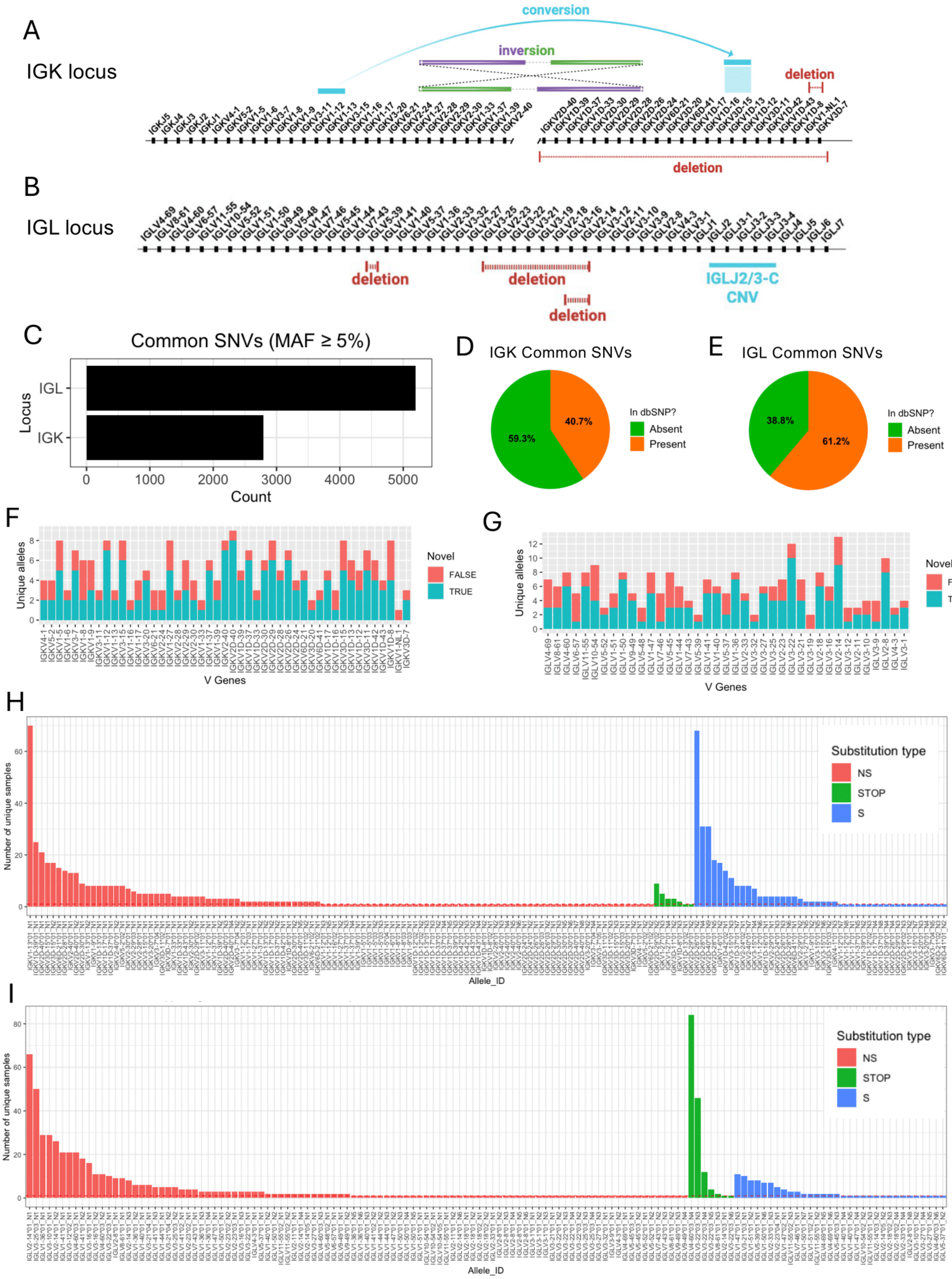
A
IGK locus
conversion
inversion
deletion
deletion
B
IGL locus
deletion
deletion
deletion
IGLJ2/3-C
CNV
C
Common SNVs (MAF ≥ 5%)
Locus
IGL
IGK
Count
D
IGK Common SNVs
In dbSNP?
Absent
Present
59.3%
40.7%
E
IGL Common SNVs
38.8%
61.2%
F
Unique alleles
V Genes
Novel
FALSE
TRUE
G
H
Number of unique samples
Allele_ID
Substitution type
NS
STOP
S
I

**Figure S2. IGK and IGL genetic variation identified by long-read sequencing in a cohort of 172 individuals.** (**A**) Diagram of genes and structural polymorphisms in the human IGK **(A)** and IGL **(B)** loci. SVs include deletions, an inversion, a CNV region in IGL (Gibson et al., 2023), as well as a region of high SNV density that likely resulted from a gene conversion event in which at least ~16 Kbp of sequence containing IGKV1-12 and IGKV1-13 replaced paralogous sequence in the distal region, described previously (Watson et al., 2015; Engelbrecht et al., 2024). (**C**) Barplot of the number of common SNVs (MAF ≥ 5%) identified in this cohort. (**D-E**) Pie charts indicate the proportion of common SNVs present and absent in dbSNP. (**F-G**) Stacked bar plots of the number of unique novel and previously curated (in IMGT) alleles for IGKV and IGLV genes. (**H-I**) Number of samples (y-axis) wherein each novel allele (x-axis, colored by substitution type) was identified. Novel alleles were first filtered to include only those with >10 supporting HiFi reads. (NS: non-synonymous, S: synonymous, STOP: premature stop codon (nonsense)).

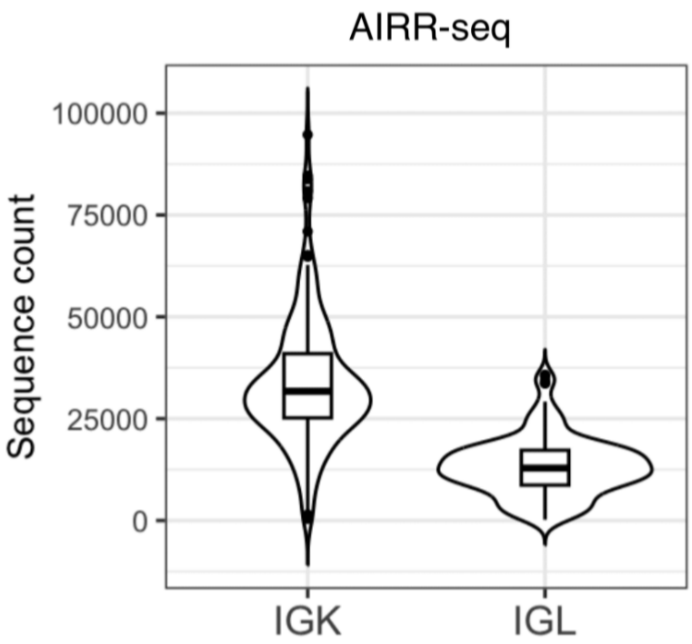


**Figure S3. Number of merged reads for each AIRR-seq dataset after processing.**
Number of unique V-J sequences per-sample after filtering duplicate reads for IGK (n=173) and IGL (n=171) AIRR-seq. Boxplots display the median, 25th percentile, 75th percentile, and whiskers that extend up to 1.5 times the inter-quartile range (IQR) from the respective percentiles. Any data points outside the whiskers are also plotted.

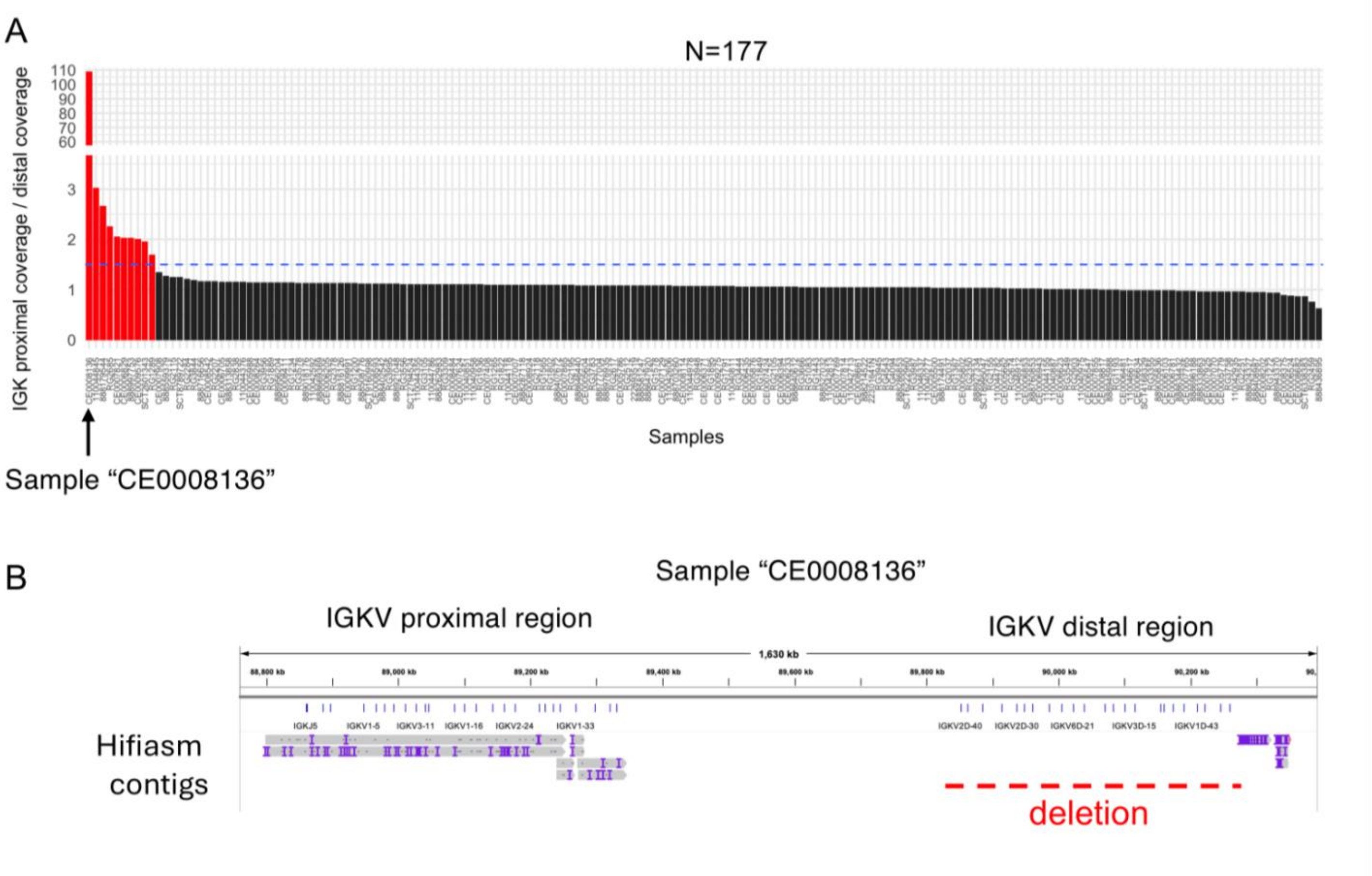


**Figure S4. Identification of putative IGKV distal region deletion by coverage analysis**
(**A**) Ratio of proximal:distal region read coverage is shown. If IGKV distal region gDNA was absent from the sample due to deletion, we anticipate this would be reflected in the computed ratio as ~2-fold more reads mapping to the proximal region as compared to the distal region. The dashed black line is at y=1.5. (**B**) IGV screenshot of hifiasm-generated contigs for sample CE0008136 aligned to the IGK locus; no contigs map to the IGKV distal region.

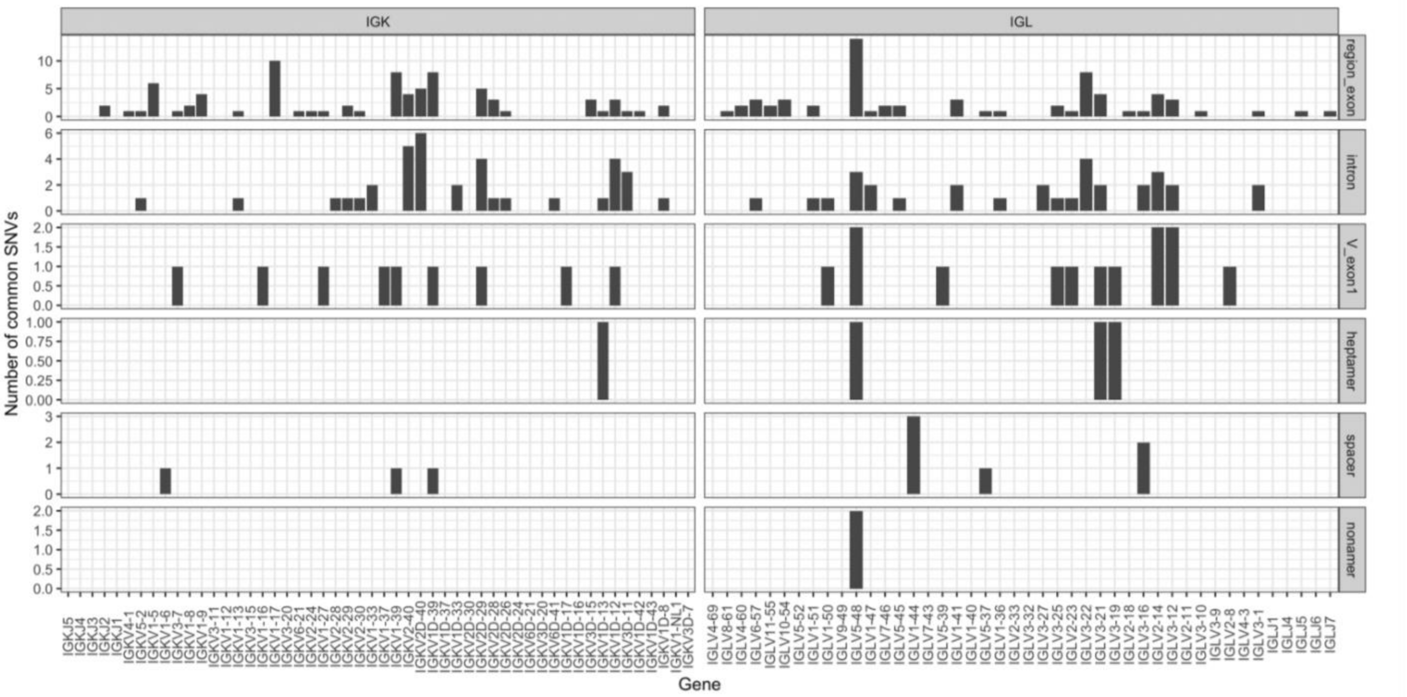


**Figure S5. Common SNVs in IGK and IGL in genic and RSS regions.**

Number of common SNVs in IGK and IGL partitioned according to location in genic and RSS (heptamer, spacer, nonamer) regions.

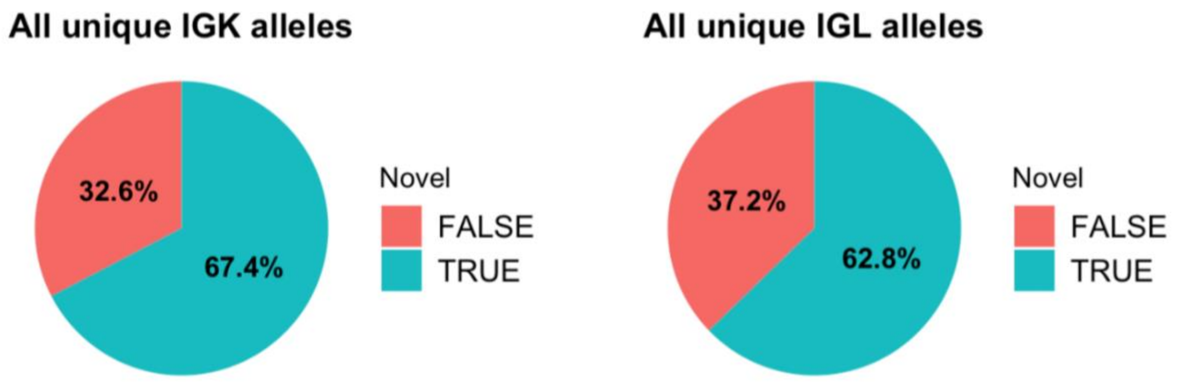


**Figure S6. Many IGK and IGL gene alleles are not catalogued in IMGT.**
Pie charts indicate the proportion of IGK and IGL gene alleles identified in our cohort that are novel (not cataloged in IMGT).

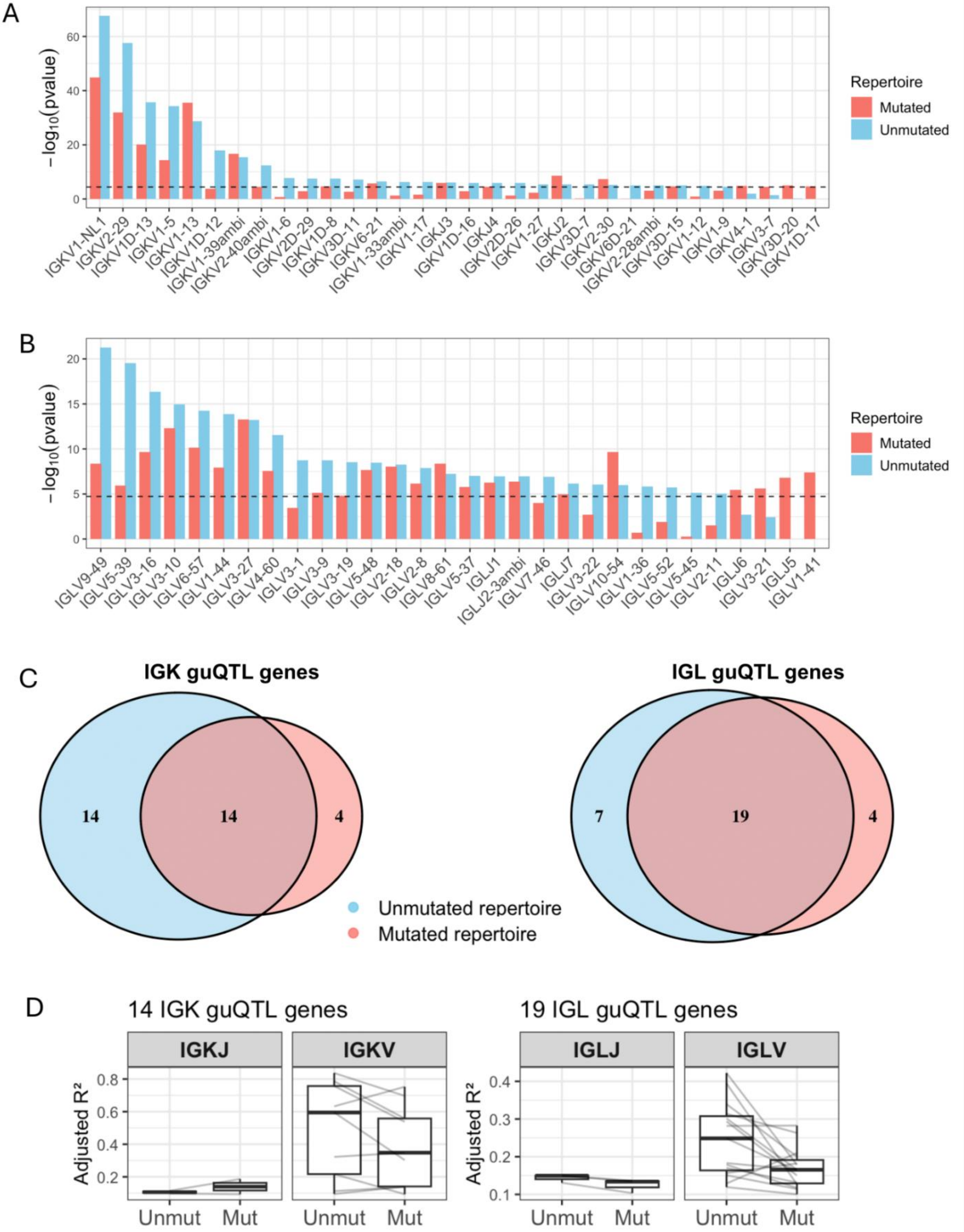
A
−log10(pvalue)
Repertoire
Mutated
Unmutated
IGKV1-NL1
IGKV2-29
IGKV1D-13
IGKV1-5
IGKV1-13
IGKV1D-12
IGKV1-39ambi
IGKV2-40ambi
IGKV1-6
IGKV2D-29
IGKV1D-8
IGKV3D-11
IGKV6-21
IGKV1-33ambi
IGKV1-17
IGKJ3
IGKV1D-16
IGKJ4
IGKV2D-26
IGKV1-27
IGKJ2
IGKV3D-7
IGKV2-30
IGKV6D-21
IGKV2-28ambi
IGKV3D-15
IGKV1-12
IGKV1-9
IGKV4-1
IGKV3-7
IGKV3D-20
IGKV1D-17
B
−log10(pvalue)
Repertoire
Mutated
Unmutated
IGLV9-49
IGLV5-39
IGLV3-16
IGLV3-10
IGLV6-57
IGLV1-44
IGLV3-27
IGLV4-60
IGLV3-1
IGLV3-9
IGLV3-19
IGLV5-48
IGLV2-18
IGLV2-8
IGLV8-61
IGLV5-37
IGLJ1
IGLJ2-3ambi
IGLV7-46
IGLJ7
IGLV3-22
IGLV10-54
IGLV1-36
IGLV5-52
IGLV5-45
IGLV2-11
IGLJ6
IGLV3-21
IGLJ5
IGLV1-41
C
IGK guQTL genes
14
14
4
IGL guQTL genes
7
19
4
Unmutated repertoire
Mutated repertoire
D
14 IGK guQTL genes
IGKJ
IGKV
Adjusted R²
Unmut
Mut
19 IGL guQTL genes
IGLJ
IGLV
Adjusted R²
Unmut
Mut

**Figure S7. guQTLs in naïve (unmutated) and antigen-experienced (mutated) Ab repertoires.**
(A-B) Barplots show the strength of associations for lead guQTLs in the unmutated (naïve) and mutated (antigen-experienced) IGK (A) and IGL (B) repertoires. Plots include only guQTL genes. (C) Venn diagrams indicating the number of unique and overlapping guQTL genes in unmutated and mutated IGK and IGL Ab repertoires (see also **Supplementary Table S7**). (D) The variance explained (adjusted $R^2$) for guQTL genes identified in both unmutated and mutated Ab repertoires for IGK (14 genes) and IGL (19 genes). Gray lines connect the variance explained in the unmutated and mutated repertoires.

Mutated repertoires

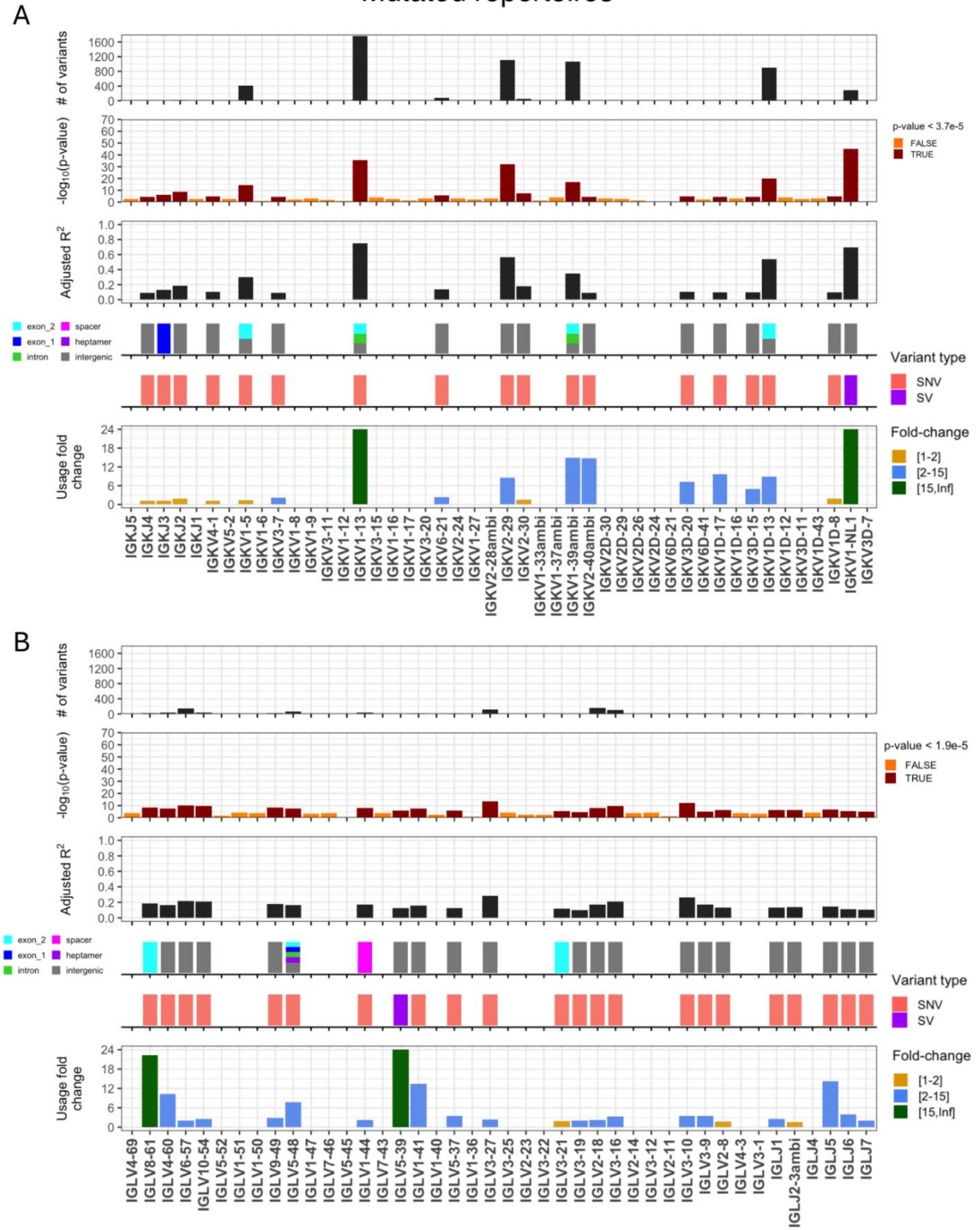


328

**Figure S8. guQTLs in naïve (unmutated) and antigen-experienced (mutated) Ab repertoires.**

(A-B) Per gene (x axis, all panels) statistics from linear regression guQTL analysis for the repertoire of mutated IGK (A) and IGL (B) light chains, including: (i) the number of associated variants after Bonferroni correction (IGK; $P < 3.7e-5$, IGL; $P < 1.9e-5$), (ii) −log10(P value) of the lead guQTL, (iii) adjusted $R^2$ for variance in gene usage explained by the lead guQTL, (iv) the location and (v) type of variant for the lead guQTL and (vi) the fold change in gene usage between genotypes at the lead guQTL. Summary statistics are provided in **Supplementary Table S6**.

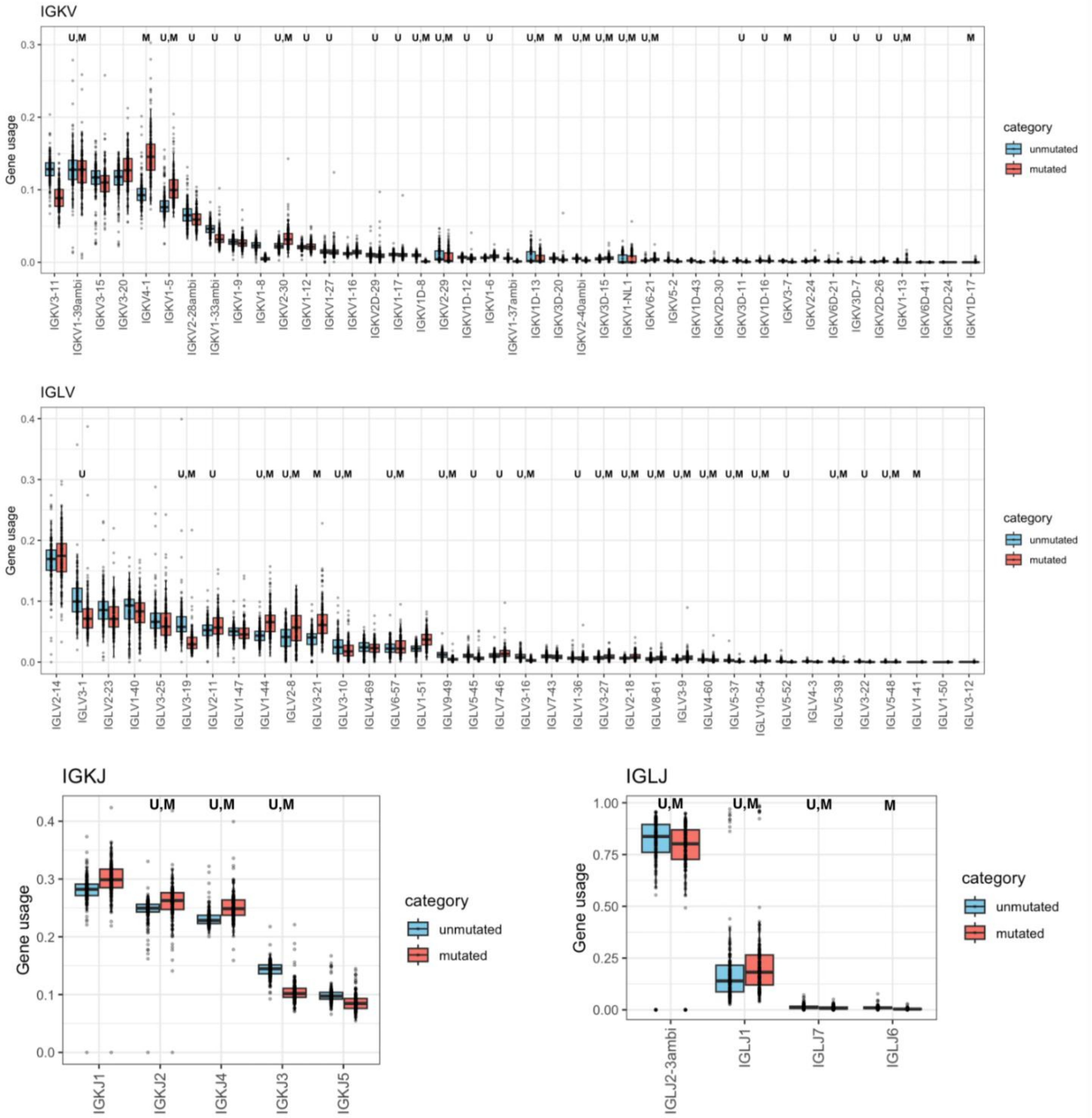


**Figure S9. Usage of IGK and IGL guQTL and non-guQTL genes in unmutated and mutated Ab repertoires.** Each panel shows usage of V or J genes in IGK or IGL. guQTL genes in unmutated and mutated repertoires are annotated "U" and "M", respectively, with "U,M" indicating guQTL genes in both unmutated and mutated repertoires.

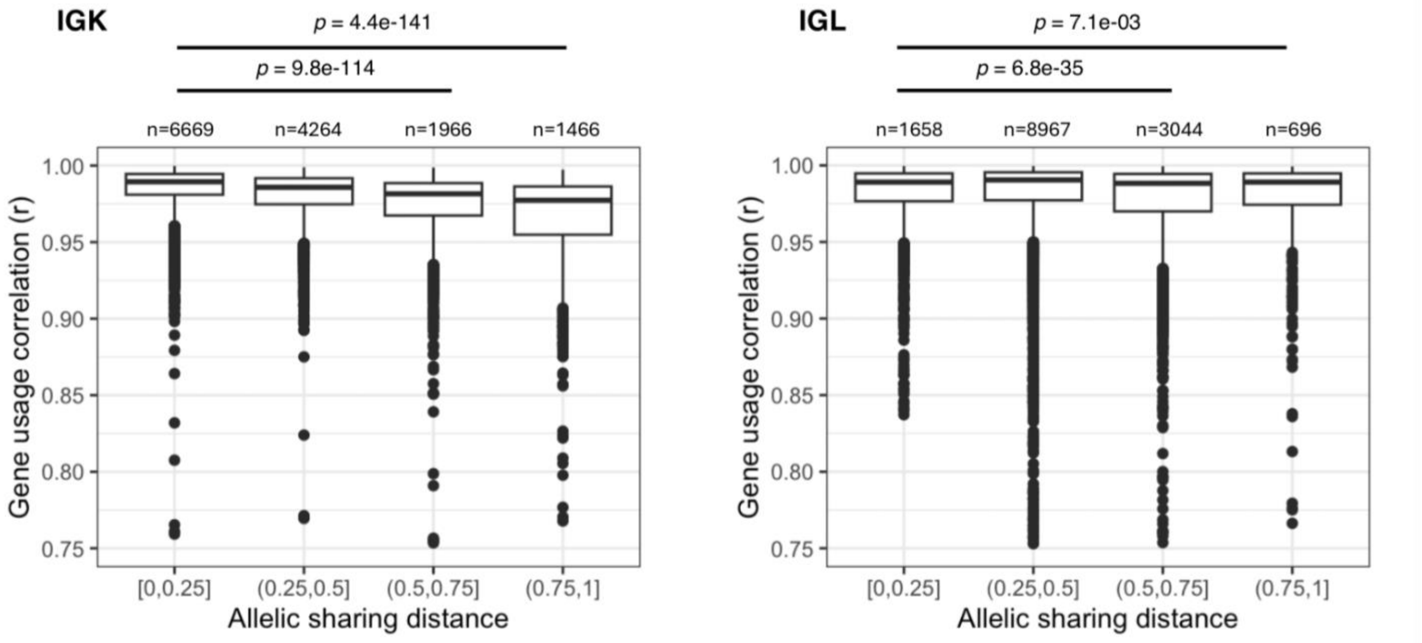


**Figure S10. Individuals sharing a greater number of guQTL genotypes have more correlated repertoire-wide light chain gene usage profiles.**

Repertoire-wide gene usage correlations between all individuals in the cohort (pairwise) were calculated using the Pearson's Correlation coefficient (y-axis). Pairs of samples are separated according to allele sharing distance (ASD), calculated using all guQTL variants in the respective locus. Boxplots plots show pairwise IGK and IGL repertoire-wide gene usage correlations partitioned by ASD. Boxplots display the median, 25th percentile, 75th percentile, and whiskers that extend up to 1.5 times the inter-quartile range (IQR) from the respective percentiles. Data points outside the whiskers are also plotted.

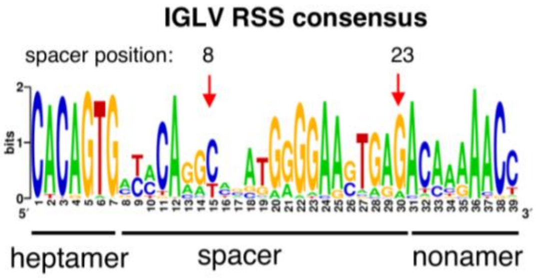


**Figure S11. IGLV RSS consensus.**

IGLV RSS consensus sequence logo computed using all unique IGLV RSSs in our cohort.

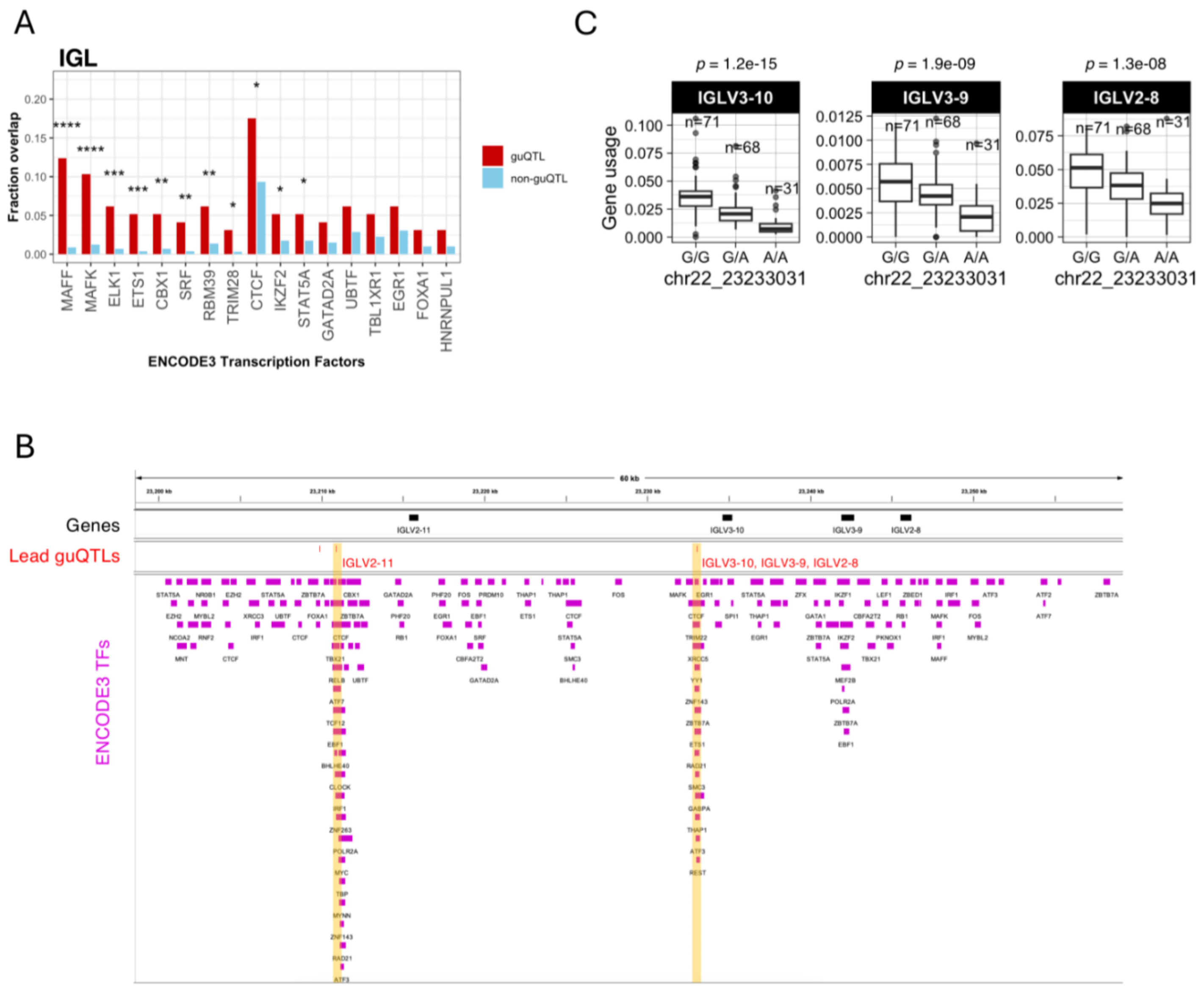


**Figure S12. IGL guQTLs are enriched with ENCODE3 TFBS.**
(**A**) Bar plot showing the fraction of lead IGL guQTL SNVs that overlapped ENCODE3 TFBS, compared to the overlap observed for the non-guQTL set of variants used in the IGL guQTL analysis. TFBS for which statistically significant enrichments were observed are indicated by asterisks: One-side Fisher's Exact Test; *P value < 0.05; **P value < 0.005; ***P value < 0.0005; ****P value < 0.00005 (see also **Supplementary Table S8**). (B) IGV screenshot with annotations for genes, guQTLs, and ENCODE3 TF binding regions. The lead guQTL for IGLV2-11 overlaps multiple TFBS. The lead guQTL for 3 IGLV genes (IGLV3-10, IGLV3-9, IGLV2-8) also overlaps multiple TFBS. (**C**) Boxplots of usages of IGLV3-10, IGLV3-9, IGLV2-8 with individuals separated by genotype at the lead guQTL variant (indicated in (**B**) for these three genes.

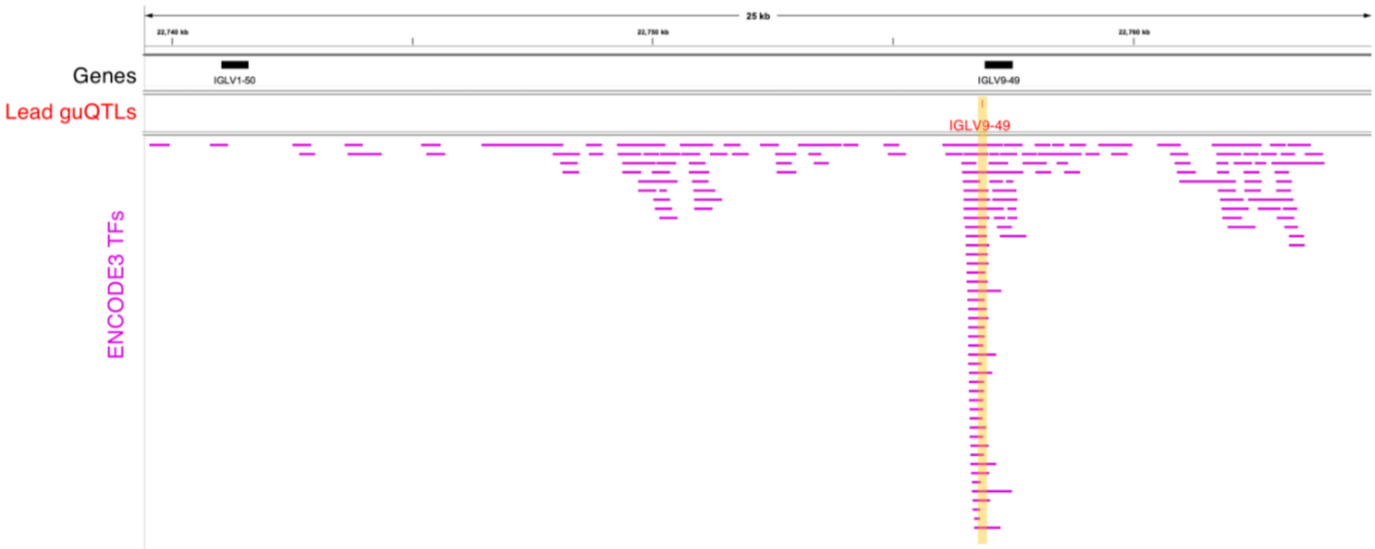


**Figure S13. The lead IGLV9-49 guQTL overlaps multiple TFBS.**

(**A**) IGV screenshot with annotations for genes, guQTLs, and ENCODE3 TF binding regions. The lead guQTL for IGLV9-49 overlaps multiple TFBS.

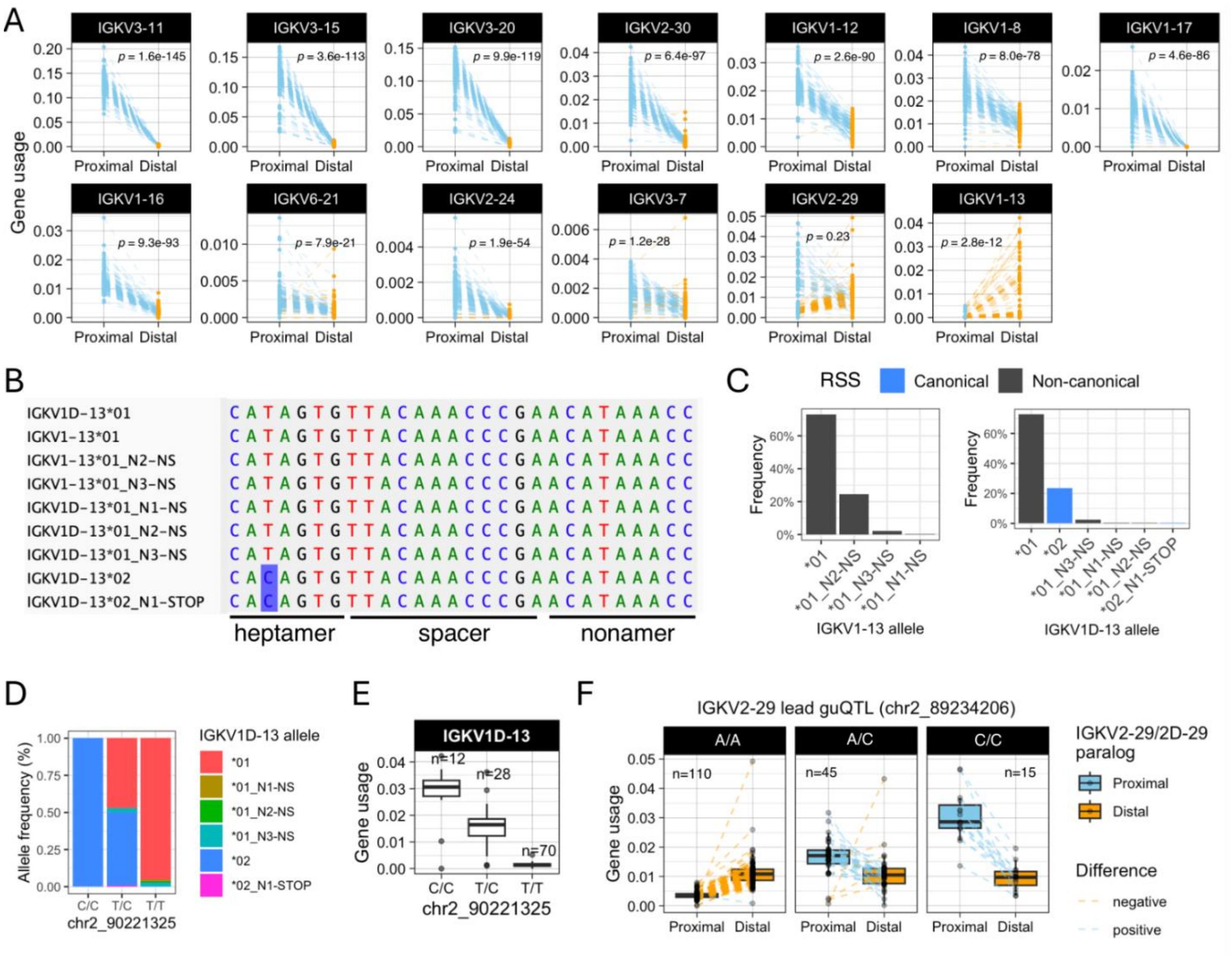


**Figure S14. Analysis of IGKV gene paralog usages.**

(**A**) Usage of indicated IGKV gene paralogs within individual (data points), with blue and orange dashed lines indicating higher usage of the proximal and distal paralog, respectively. Differences between proximal and distal gene usage were determined by paired t-tests. (**B**) Alignment of all RSSs for IGKV1-13 and IGKV1D-13 alleles in our cohort. (C) The frequency of each IGKV1-13 and IGKV1D-13 allele in our cohort, with alleles colored according to having a canonical or non-canonical RSS heptamer. (D) The frequency of IGKV1D-13 alleles in lead guQTL IGKV1D-13 genotype groups. (E) Boxplot of IGKV1D-13 usage with individuals separated according to genotype at the lead IGKV1D-13 guQTL. (F) Boxplot of IGKV2-29 (proximal) and IGKV2D-29 (distal) gene usages in individuals, with blue and orange dashed lines indicating higher usage of the proximal and distal paralog, respectively. Individuals are grouped (columns) according to genotype at the lead IGKV2-29 guQTL (discussed in **Figure 2A-C**).

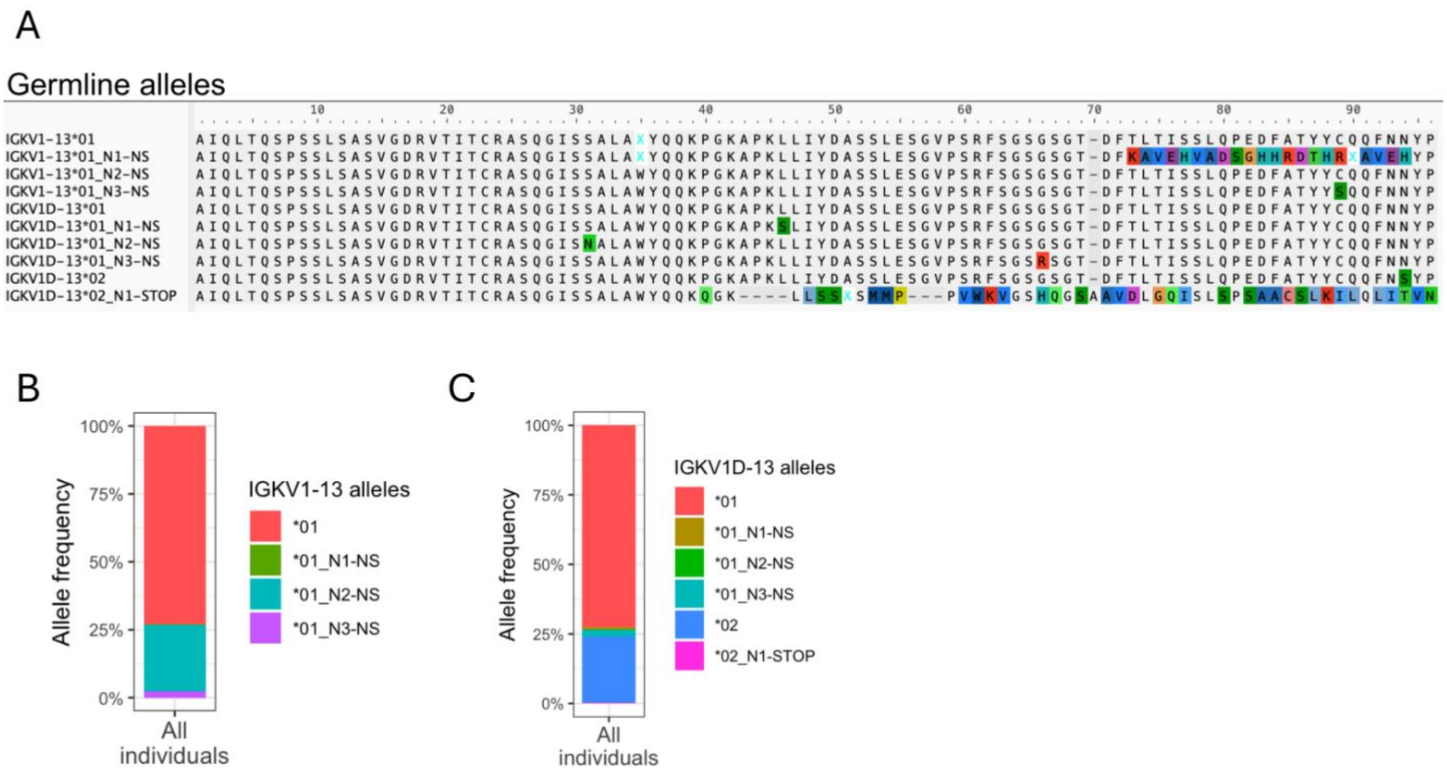


**Figure S15. IGKV1-13 and IGKV1D-13 germline alleles.**

(**A**) Alignment of translated V-regions of IGKV1-13 and IGKV1D-13 alleles. (**B-C**) Frequency of IGKV1-13 (B) and IGKV1D-13 (**C**) alleles among individuals in the cohort.

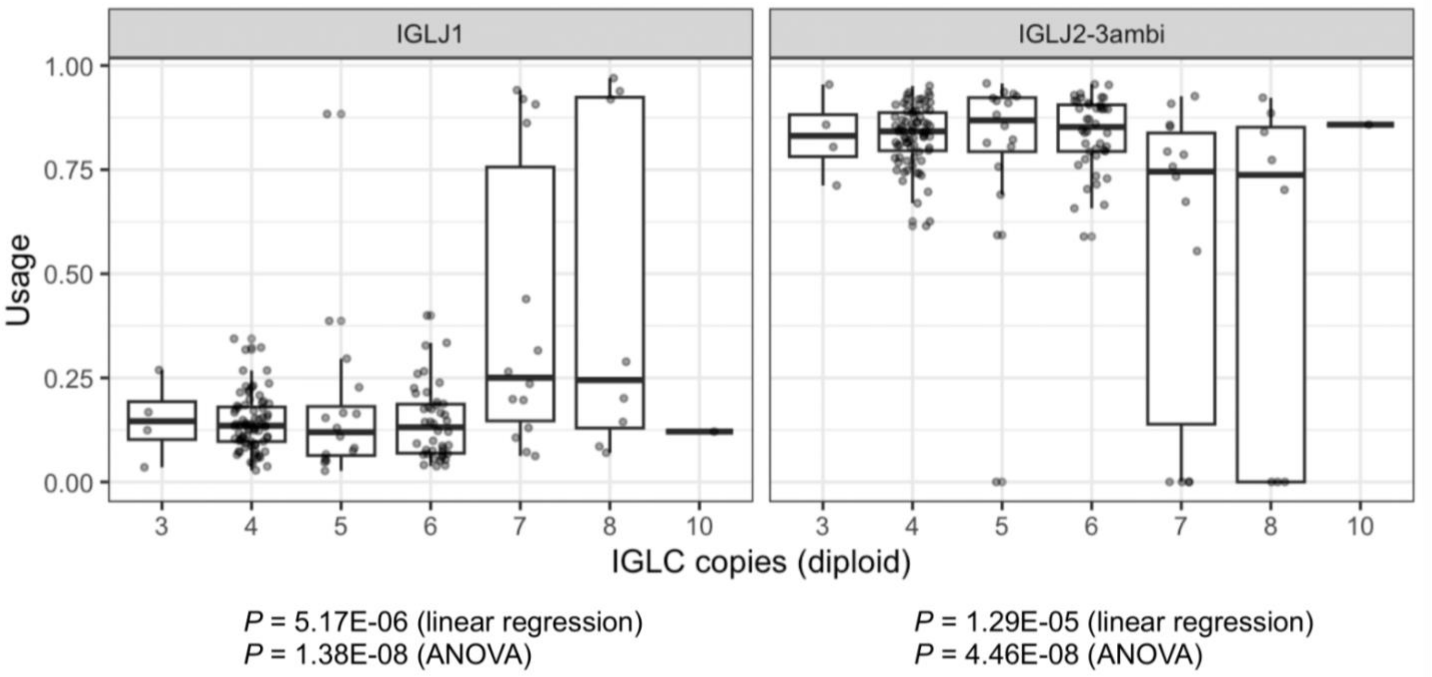


**Figure S16.** Usage of IGLJ1 and IGLJ2-3ambi, with individuals grouped according to number of diploid copies of the IGLJ2-3 cassette. P values from indicated tests are shown.

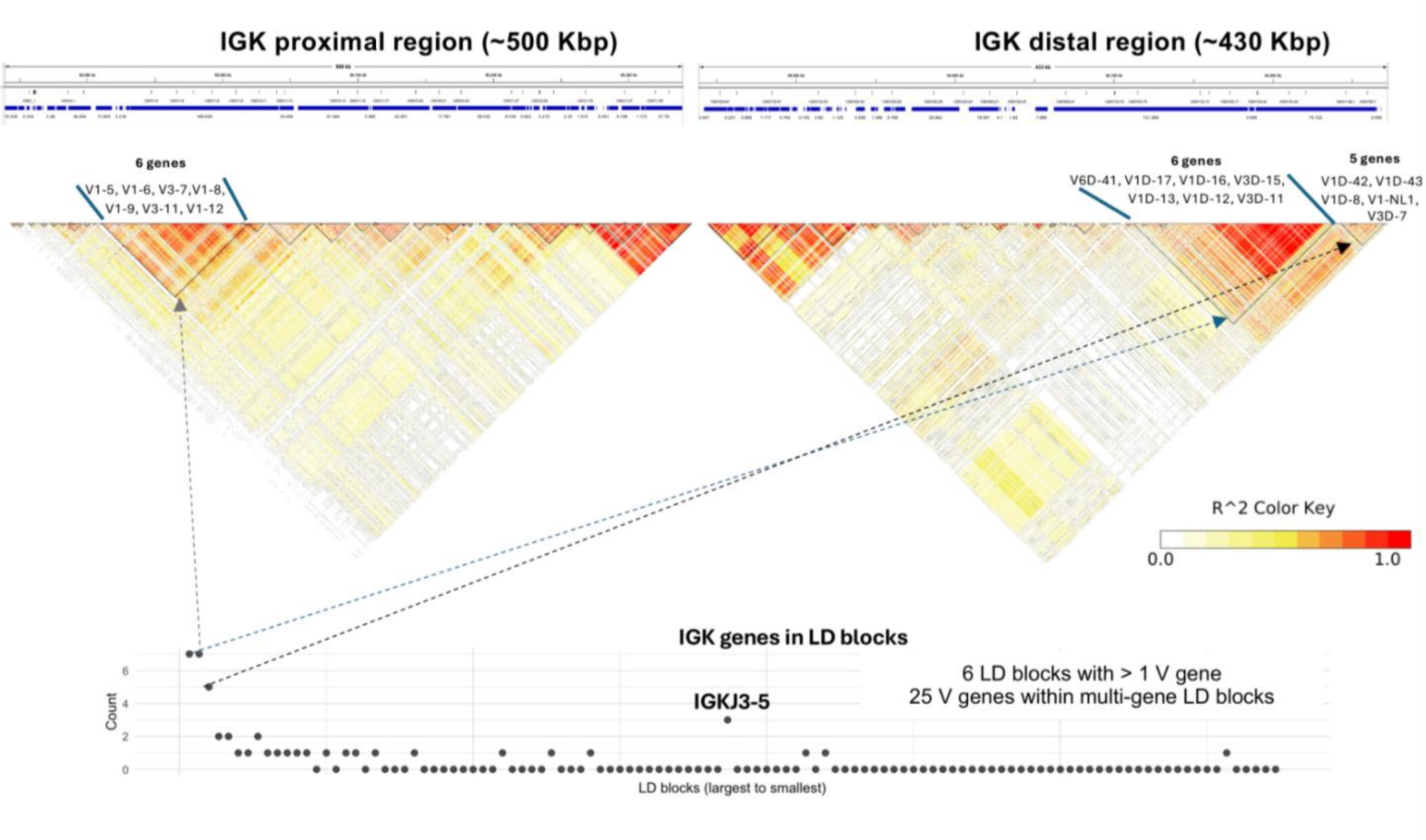


**Figure S17. LD blocks in IGK.**

(Top) Heat corresponds to pairwise correlations between SNVs; LD blocks (black triangles) were computed using LDBlockShow (Dong et al., 2021). (Bottom) LD blocks are plotted along the x-axis from largest to smallest (left to right), and the count of the number of genes within each block is shown. Arrows are drawn for the 3 largest LD blocks.

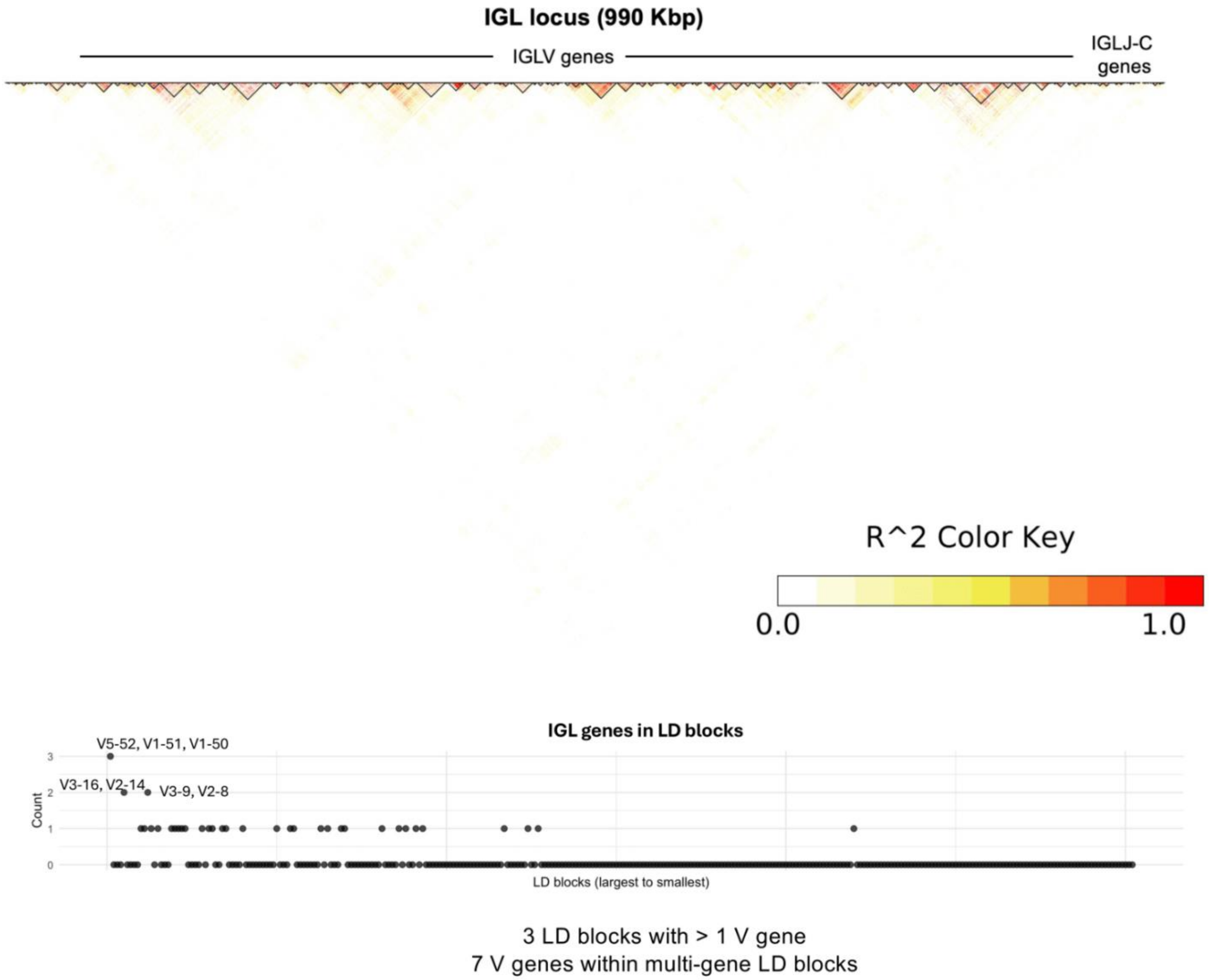


**Figure S18. LD blocks in IGK.**

(Top) Heat corresponds to pairwise correlations between SNPs; LD blocks (black triangles) were computed using LDBlockShow (Dong et al., 2021). (Bottom) LD blocks are plotted along the x-axis from largest to smallest (left to right), and the count of the number of genes within each block is shown. Genes within the 3 largest LD blocks are indicated.

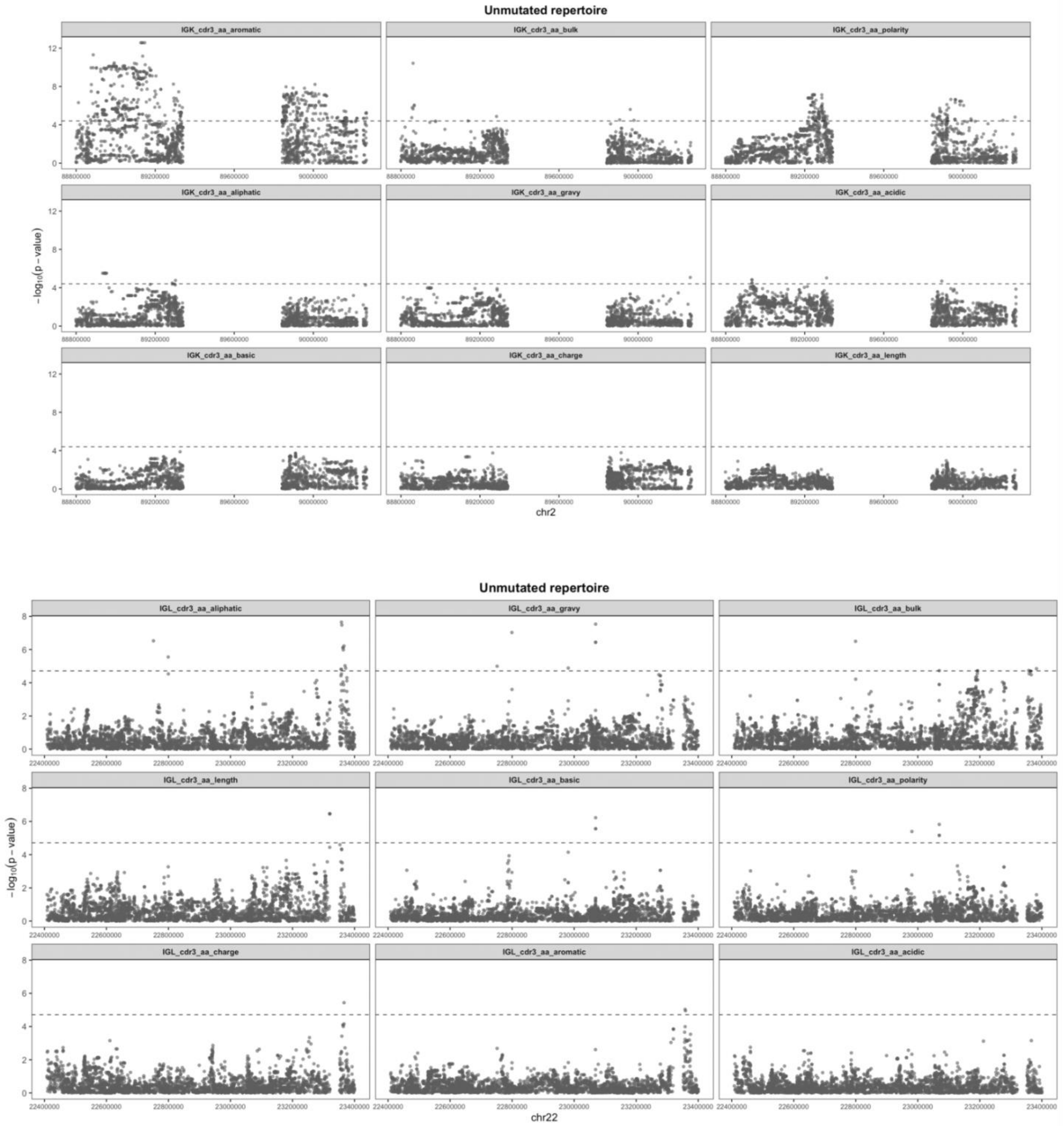


**Figure S19. Associations between CDR3 physicochemical properties and germline variants in IGK and IGL.** Manhattan plots show the −log10(P value) for all SNVs in IGK (top) or IGL (bottom) tested for association with indicated CDR3 physicochemical properties in naïve (unmutated) Ab repertoires. Dashed lines indicate Bonferroni-corrected significance (IGK; $P < 3.7e\text{-}5$, IGL; $P < 1.9e\text{-}5$).

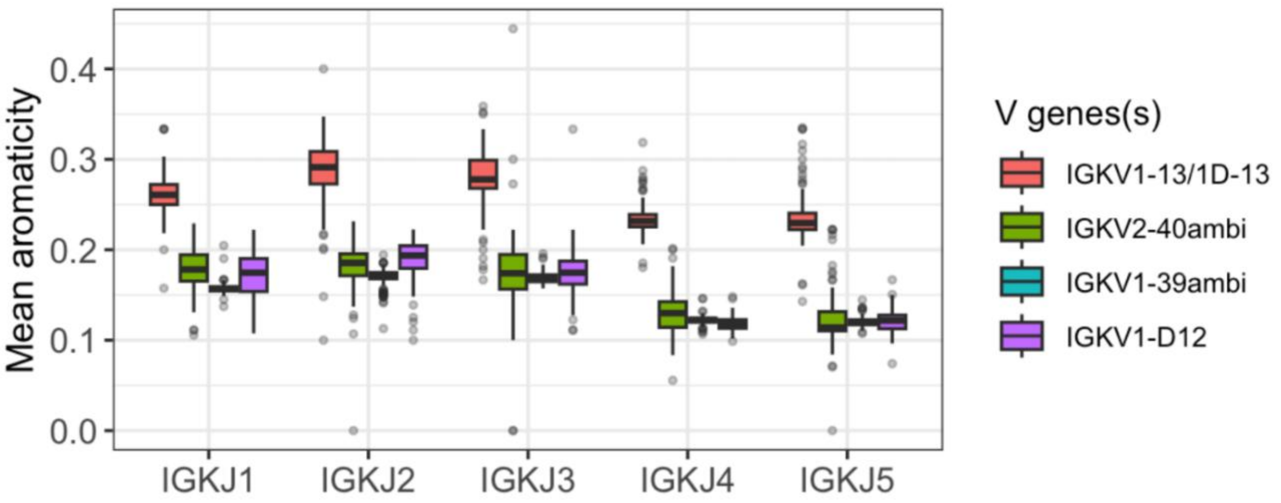


**Figure S20. CDR3 aromaticity of BCR sequences composed of specific IGKV and IGKJ genes.**

Boxplot of mean CDR3 aromaticity (per-individual) of unmutated IGK BCR sequences comprised of indicated IGKV and IGKJ gene pairs.

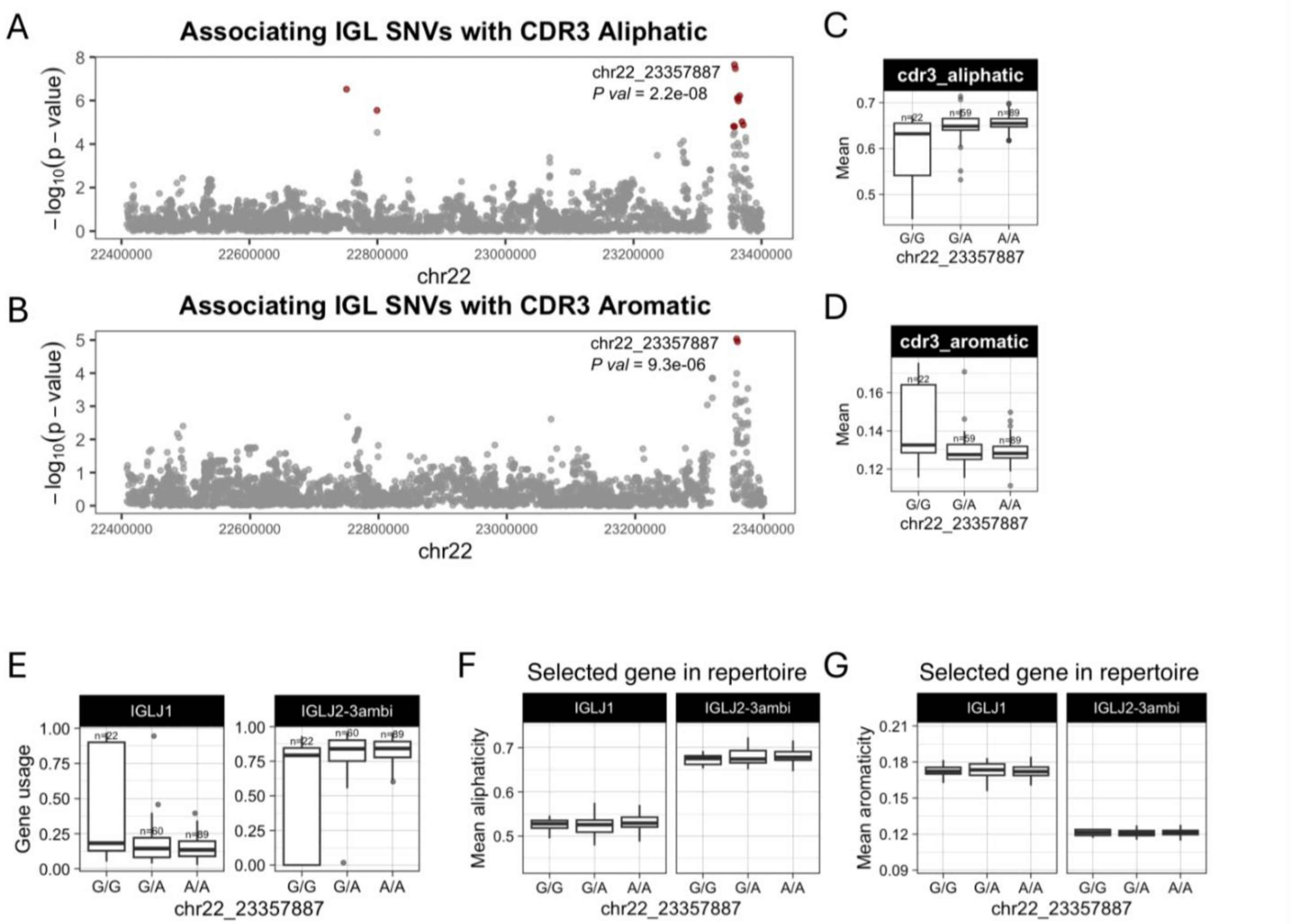


**Figure S21. Genetic effects on IGLJ gene usages are associated with CDR3 aliphaticity and aromaticity.** (**A-B**) Manhattan plots shows the −log10(P value) for all SNVs in the IGL locus tested for association with CDR3 aliphaticity (**A**) or aromaticity (**B**), with QTLs colored dark red and the lead QTL labelled. These two CDR3 properties share a lead QTL variant (labelled). (**C-D**) Boxplots of the mean IGL CDR3 aliphaticity (**C**) and aromaticity (**D**) with individuals separated by genotype at the lead QTL. (**E**) Boxplot of IGLJ1 and IGLJ2-3ambi usages with individuals separated by genotype at the lead guQTL. (**F-G**) BCR sequences that used IGLJ1 or IGLJ2-ambi were selected from the Ab repertoire, then mean CDR3 aromaticity of each repertoire subset was computed and plotted with individuals separated by genotype at the lead variant.
